# Deep learning to overcome human error and bias in electrode position extraction in tDCS-fMRI studies

**DOI:** 10.64898/2025.12.23.696193

**Authors:** F. Niemann, S. Riemann, S. Dabelstein, K. Hering, H. Kocataş, M. Abdelmotaleb, L.M. Caisachana Guevara, R. Fischer, A. Thielscher, D. Antonenko, A. Flöel, M. Meinzer

## Abstract

Combining transcranial direct current stimulation (tDCS) with fMRI enables investigation of stimulation effects, while structural MRI verifies electrode positioning, critical for focal montages where minor misplacements reduce target current dose. Currently, no fully automated methods exist for precise electrode extraction from MRI.

We developed an Attention U-Net-based deep learning approach to automate electrode detection in focal tDCS-fMRI studies, using structural pointwise-encoding time reduction with radial acquisition (PETRA) MRI scans from a multicenter trial (N = 392 images; 1,568 electrodes, https://www.memoslap.de/en/home/). Performance was compared to manual and semi-automated methods for a 3x1 montage (three cathodes around a central anode). The network achieved robust segmentation (Dice Score = 0.76, Hausdorff distance = 36.76 mm), identifying all electrodes in 95% of cases (323/340). These metrics compared network-segmented electrodes to highly accurate, manually segmented electrodes, which are also called "ground truth". Linear mixed-effects models on 52 “ground truth” images showed deep learning outperformed manual and semi-automated methods, aligning best with ground truth. Fully/semi-automated methods comparison of 290 images showed highest agreement (ICC = 0.990, bias = 1.3 mm), while manual extraction exhibited larger biases (−5.77 to −7.06 mm) and systematic errors.

The automated approach overcomes manual limitations by improving precision, eliminating human variability and bias, and enabling scalability for large studies. It sets a new standard for electrode verification in focal tDCS, particularly for studies requiring precise localization. The model is open-source, with future refinements discussed

## 1. Introduction

Transcranial direct current stimulation (tDCS) has widely been used to modulate human brain functions in research and clinical settings (Chan et al., 2021; Kang et al., 2024; Narmashiri and Akbari, 2025; Perceval et al., 2016; Simonsmeier et al., 2018). By applying weak electrical currents via scalp-attached electrodes, tDCS induces shifts in neuronal membrane potentials, resulting in transient changes of cortical excitability (Stagg et al., 2018) Combining functional magnetic resonance imaging (fMRI) with concurrently applied tDCS has emerged as a key tool for investigating these acute effects of tDCS on large-scale brain networks (Esmaeilpour et al., 2020; Meinzer et al., 2024) This approach also allows verification of electrode placement accuracy on the scalp by using structural imaging data acquired during the same imaging session (Meinzer et al., 2014; Niemann et al., 2024b). The latter is relevant, because deviations of electrodes from intended scalp positions due to mis- or displacement, can affect the distribution and intensity of the induced electric field in the brain (Indahlastari et al., 2023; Woods et al., 2015). Focal tDCS set-ups that constrain the current flow to circumscribed brain regions (Niemann et al., 2024b) are particularly affected by such errors. For example, positioning errors of 5 mm in a focal motor cortex set-up reduced peak somatic polarization in the hand knob by up to 87% (Seo et al., 2017). Moreover, a recent computational study showed that electrode positioning errors resulted in significantly more pronounced dose reductions (26-43%) in the target regions for focal compared to conventional set-ups (Niemann et al., 2024a). This highlights the relevance of verifying electrode positioning accuracy during intra-scanner tDCS-fMRI studies and incorporation of actual electrode positions in computational models of current flow involving focal set-ups, for example when investigating dose-response relationships (Ekhtiari et al., 2022; Meinzer et al., 2024). However, there are currently no fully automated methods available to accurately, reliably and efficiently determine tDCS electrode positions from structural MRI.

So far, the vast majority of tDCS-fMRI studies have either not verified electrode positions (Hampstead et al., 2020; Leaver et al., 2023; Müller et al., 2023; Rauh et al., 2023; for reviews see Esmaeilpour et al., 2020; Ekhtiari et al., 2022) or relied on visual inspection, manual segmentation or semi-automatic extraction from 3D or 3D-to-2D converted images (e.g., Antonenko et al., 2019; Indahlastari et al., 2021, 2023; Niemann et al., 2024a). All approaches are time consuming and subject to human error. Alternative semi-automated options, originally developed for identifying EEG electrode positions, require specific equipment (i.e., 3D digitizers, artificial markers or laser scanners, infrared optic stereo tracking systems) and transformations of electrode positions from pointer devices to MRI space can introduce inaccuracies (Nielsen et al., 2023; Sijbers et al., 2000). A semi-automatic method proposed by De Munck et al.(2012) addressed these challenges by utilizing the artifacts left by electrode gel on anatomical MRI scans, providing a more direct and efficient approach.

In the present study, we modified this approach and integrated it into a deep learning framework for fully automated tDCS electrode coordinate extraction from structural pointwise-encoding time reduction with radial acquisition (PETRA) MRI scans. Deep learning is a machine learning approach that uses multilayered artificial neural networks that excel in classification tasks, like image segmentation (Goodfellow et al., 2016; LeCun et al., 2015). Networks typically employ supervised learning, requiring labeled training data (the *ground truth*) to learn how to identify relevant patterns (e.g., electrodes on the scalp), prior to attempting extraction from previously unseen images.

The new fully automatic approach we developed was based on an U-Net architecture (i.e., a form of convolutional neural networks), widely used for medical image segmentation (Ronneberger et al., 2015) due to their encoder-decoder architecture, with skip connections to preserve spatial details, while capturing contextual information. For small structures in MRI (e.g., electrodes, lesions, or subcortical nuclei), standard U-Nets may struggle with fine-scale features due to information loss in deeper layers. Attention Gates (AGs) enhance U-Nets by dynamically weighting so-called “skip connections” that focus the model on relevant regions, while suppressing background noise (Lim et al., 2021; Oktay et al., 2018; Vaidyanathan et al., 2021). For training and validation of the network, we provided highly accurate, manually segmented and labeled electrodes from an ongoing large-scale tDCS-fMRI study as “the ground truth”. Performance was subsequently evaluated using a separate dataset, acquired at the same scanner with identical tDCS montages. A previous feasibility study (Niemann et al., 2025) showed that our network performed similarly to a semi-automated method, also developed by our group (Niemann et al., 2024b). However, that analysis did not distinguish between ground truth and extended data, nor did it analyze manual segmentation errors. In this current work, we compared the outcomes of our new approach to both manual and semi-automated methods for tDCS electrode position extraction, which were previously described by our group (Niemann et al., 2024a, 2024b). We also performed a separate analysis of the 54 ground truth images from the extended dataset of 340 images not seen by the network.

Additionally, we have included the most up-to-date segmentation results from 1,823 images (including 72 ground truth data) in the limitations section to demonstrate the network’s generalization ability.

## 2. Material and Methods

### 2.1 Study overview

To evaluate performance of our deep learning-approach for fully automated electrode coordinate extraction, we used data collected in three projects of an ongoing collaborative research consortium (Research Unit 5429, https://www.memoslap.de, accessed July 29.2025). The consortium employs concurrent tDCS-fMRI to investigate modulation of learning and memory by focalized tDCS across different functional domains. All projects use the same 3x1 set-up (i.e., a central anode and three surround cathodes) to administer focalized tDCS to the left temporoparietal junction (lTPJ), right dorsolateral prefrontal cortex (rDLPFC) or right occipito-temporal cortex (rOTC). Montages were optimized for each study participant by individualized current modeling (based on structural MRI data acquired during a baseline session and positioning of electrodes on the scalp was guided by neuronavigation (Niemann et al., 2024b).

Electrode configurations (i.e., the distance between anode and the three cathodes and between the cathodes) were standardized using a 3D-printed spacer (Niemann et al., 2024a). Participants completed up to four resting-state or task-based fMRI sessions with concurrent tDCS. Structural pointwise-encoding time reduction with radial acquisition (PETRA; TR = 3.3ms, TE= 0.07, Slice per slab = 320, Slice thickness = 0.9) scans were acquired before and after the respective functional sequences and used for electrode position verification. The sharp contrast between the gel and the electrode makes PETRA scans without fat suppression particularly suited for the automated segmentation task intended here (Grodzki et al., 2012). All data were acquired at the same Siemens MAGNETOM Vida 3T scanner at the University Medicine Greifswald using a 64-channel head/neck receive coil and the syngo_MR_XA50 software. The study was approved on February 20, 2020, by the local ethics committee (reference number BB 172/19) and was conducted in accordance with the Helsinki Declaration. Written informed consent was obtained prior to inclusion in the study.

### 2.2 Data

All participants completed a baseline (f)MRI session (including PETRA scans without electrodes attached). PETRA scans acquired during subsequent fMRI sessions (1-4) and runs (i.e., pre-post fMRI) were co-registered onto the first baseline PETRA scan using Statistical Parametric Mapping software (SPM12, https://www.fil.ion.ucl.ac.uk/spm/, accessed July 29.2025). In total we had access to 392 images from 57 participants (27 men, 29 women, 1 non-binary, 25.05 ± 5.46 years). For manual (**Figure 1A**) and semi-automatic (**Figure 1B**) data extraction, we included a total of 363 PETRA images with electrodes attached from 48 participants (22 men, 25 women, 1 non binary, M ± SD age: 25.09 ± 5.62 years). Twelve sessions from eight participants were unavailable (because participants did not complete 1-2 imaging sessions, due to corrupt data or data loss during transfer). The automatic network-based extraction approach ( **Figure 1C**) was conducted subsequently and initially had access to the original 392 images. 52 of those images were randomly selected within the set of 392 images and 2 additional images from later measured participants for a total of 54 images. These images were used to determine the ground truth (see below). The remaining 340 images were submitted to network-based segmentation and post-processing to verify if the correct number of electrodes (N=4) were identified.

**Figure 1:**
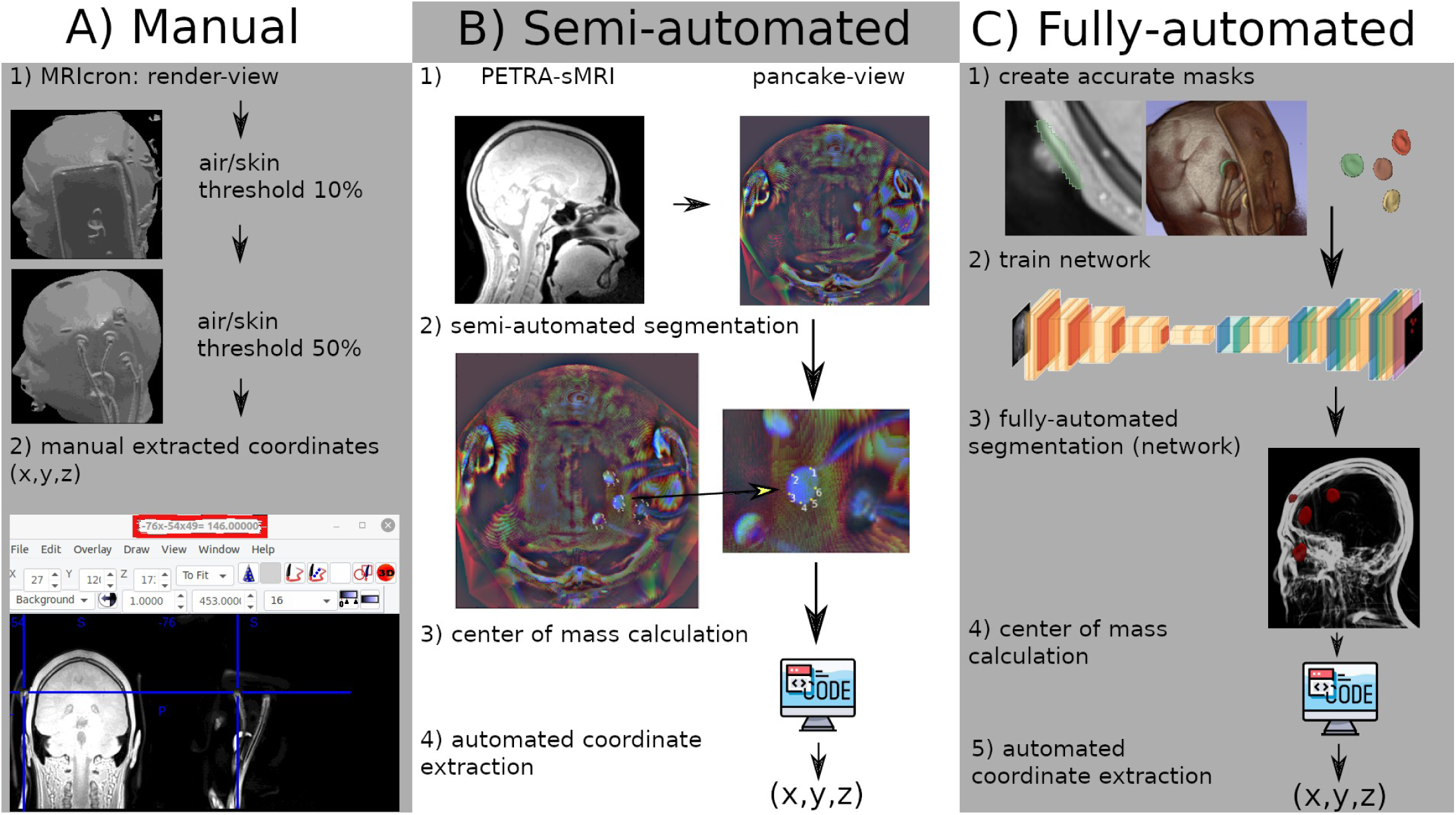
Methods overview. Methods overview of manual, semi- and fully automated electrode extraction. Schematic workflow is given for each method. PETRA-MRI (Pointwise Encoding Time Reduction with Radial Acquisition-Structural Magnetic Resonance Imaging)

### 2.3 Fully automatic (network metric) approach

#### 2.3.1 Label acquisition

Ground truth electrode masks were generated by manually segmenting 54 randomly selected images from 38 participants (lTPJ: 12 images/11 participants; rDLPFC: 14 images/12 participants; rOTC: 28 images/15 participants) using 3D Slicer (Fedorov et al., 2012; **Figure 1C1**). All four electrodes per image were labeled as ground truth (i.e., a combined ground truth mask). These 54 images were used for training, validation, and testing, enabling pixel-level segmentation between electrodes and background across different montages. By using different montages that targeted three different brain regions for training, we aimed to ensure robust electrode segmentation for broader data sets (Vrudhula et al., 2024).

#### 2.3.2 Network implementation

We employed a 3D Attention U-Net architecture (see **Figure 1 C2**, for further details see **Figure B.1**) for automated electrode segmentation from PETRA scans (see **Figure 1 C3**). The network was implemented via python based packages PyTorch (Paszke et al., 2019) and MONAI (Medical Open Network for AI; (Cardoso et al., 2022)), specially developed for deep learning approaches in medical data, on an NVIDIA RTX 5000 Ada Generation graphic card. The Attention U-Net built upon the standard U-Net framework (Buda et al., 2019), while incorporating attention mechanisms to enhance segmentation accuracy, particularly beneficial for small structures (like scalp attached electrodes). The key advantages of AGs for detecting small structures include (a) improved localization precision by highlighting salient features (e.g., electrode edges) in skip connections for better boundary delineation, (b) reduced redundancy through filtering of irrelevant activation, (c) mitigation of class imbalance issues common in small-object segmentation and (d) enhanced computational efficiency by avoiding costly full attention mechanisms.

The architecture consisted of a symmetric encoder-decoder structure with five hierarchical levels. The encoder path progressively reduced spatial resolution while increasing feature channels (32, 64, 128, 256, and 512 channels at successive levels), using stride convolutions with a stride length of 2 for down-sampling. Decoders reverse this process, i.e., resulting in increased spatial resolution and reduced feature channels. Between the encoder and decoder paths, attention gates learned to focus on relevant spatial regions while suppressing irrelevant background areas, which proved particularly beneficial for identifying small electrode structures. The network processed single-channel gray scale MRI volumes and produced two-channel output maps distinguishing between background and segmented electrodes. All convolutional operations employed 3×3×3 kernels, with transposed convolutions in the decoder path using the same kernel size. A dropout rate of 0.2 provides regularization throughout the network.

The dataset was split into training (70%), validation (15%), and test (15%) sets, ensuring proper model evaluation (Goodfellow et al., 2016; Lim et al., 2021). Training data were used during the training protocol, whereas validation end test data were used for model evaluation. MRI volumes were processed with their original resolution (224×288×288 voxels). Extensive data augmentation enhanced the model’s robustness, including random spatial transformations (zooming between 0.7- 1.3 × scale, rotating up to ±0.5 radians in all axes, affine transformations, and flipping), intensity variations (Gaussian noise, smoothing, contrast adjustment, and intensity shifting), and regularization techniques like coarse dropout. These augmentations were determined to be optimal to improve model generalization in an iterative adaptation process, while maintaining the anatomical plausibility of the transformed images.

#### 2.3.3 Training protocol

The training protocol employed a Dice-Focal loss function, i.e., a weighted average Dice score and Focal loss (λ_dice_=0.3, λ_focal_=0.7), with loss parameters set to γ=2.5, using softmax activation, one-hot encoded targets to balance the small electrode foreground and large background, and also easy and hard to segment electrodes (Zhao et al., 2020). An Adam optimizer (Kingma and Ba, 2015) drove the learning process with gradient accumulation over eight steps to effectively increase the batch size, while managing memory constraints. The model trained with a nominal batch size of 1 due to memory limitations imposed by the 3D volumes, proceeding for up to 50,000 iterations. Additional regularization came from weight decay (1e-4) and the aforementioned dropout and data augmentation strategies.

#### 2.3.4 Model evaluation

Model performance was evaluated using the Dice Similarity Coefficient (DSC) to measure volumetric overlap accuracy between ground truth and the network’s segmented voxels . One additional metrics was used for performance tracking: Hausdorff Distance (HD) for boundary agreement (for an overview of metrics used in medical image segmentation see (Taha and Hanbury, 2015). As a benchmark for tumor segmentation with an Attention U-Net, Dice similarity coefficient (DSC) scores range from 0.78 to 0.88, while Hausdorff distance (HD) values range from 26.66 to 7.79 (Yousef et al., 2023). For a comprehensive overview of DSC metrics across various U-Net architectures and target regions, see Isensee et al. (2021). Metric calculation was restricted to the image foreground and ignored empty slices to focus on relevant areas. During training, the model was periodically validated after 500 steps (or 13 epochs) on a separate set (validation set), with checkpointing to save the best-performing weights based on the validation Dice score. After training, the top-performing model was evaluated on a held-out test set for unbiased performance assessment. The final model was applied to all data not included in the network’s training process for fully automatic electrode segmentation. The center of mass was then computed to determine the electrode coordinates (see **Figure 1 C4-5**).

#### 2.3.5 Post-processing

a. To address false positive segmentations of electrodes, we implemented a post-processing filter that triggered when more than four electrodes were detected. Through visual inspection of inference results, we observed that most false positives appeared as small additional segmentation adjacent to actual electrodes, often capturing parts of the electrode not fully covered by the primary segmentation. These artifacts were systematically removed by eliminating any segmentation clusters smaller than 20% of the average cluster size across all detected electrodes. If this process did not yield four electrodes, subjects were excluded from further analysis, as this prevented the next post-processing step (i.e., labeling), which was the case for 17 images out of 340.
b. Regarding electrode naming conventions, we standardized and automated the labeling of cathodes (C1–C3) across manual, semi-automated, and fully automated segmentation. We aligned them with coordinate naming schemes derived from semi-automatic coordinate extraction. We compared the distances between the labels of the fully automated and semi-automated electrodes and changed the labels if they were not the closest. The labels were then checked for uniqueness, and images that had the same label for multiple coordinates or vice versa were excluded. Images with inconsistent electrode renaming, typically due to excessive coordinate extraction errors, were excluded from further analysis (which was the case for 33 images out of 323).

### 2.4 Manual and semi-automatic electrode extraction

For simple <u>manual electrode extraction</u>, each PETRA image was opened using the render function of MRIcron (Rorden, 2025) to generate a 3D view of the participants’ heads with electrodes attached. For each target region, a predefined Azimuth and Elevation value was used to prevent angle dependent errors (lTPJ (Azimuth=270, Elevation=30), rDLPFC (120,30), rOTC (60,0)). The air/skin threshold was adjusted until electrodes were no longer masked by inflatable cushions (used during image acquisition to stabilize the participants’ heads in the scanner, see **Figure 1 A1**). Staff performing the manual extraction were instructed to clicked as precisely as possible at the center of each electrode and to record the resulting MNI-coordinates ( **Figure 1 A2**). <u>Semi-Automatic coordinate extraction:</u> Because identification of electrode positions in 3D is prone to errors, a modified 2D “pancake” view of the reconstructed scalp with electrodes was used (see **Figure 1 B1**; De Munck et al., 2012; Fleury et al., 2019) to determine six edge points of the center anodes (see **Figure 1 B2)**. This method was adapted (Niemann et al., 2024b), because the workflow suggested by De Munck et al. (2012) did not reconstruct electrodes used for intrascanner tDCS. Subsequently, the center of mass was calculated (see **Figure 1 B3)**, to obtain the center coordinates of the electrodes for each fMRI session and time-point (i.e., PETRA images acquired before and after functional imaging). Subsequently, a reverse 2D to 3D transformation was applied to obtain electrode coordinates in 3D space (**Figure 1 B4)**. The method was implemented via the scripts from Butler (https://github.com/russellu/ute_git, Butler, 2016) and can be accessed via https://github.com/LawsOfForm/Automated_Electrode_Coordinate_Extraction (Niemann et al., 2024b, accessed July 29.2025). Inter-rater reliability was assessed for both manual (**Figure A.1**) and semi-automatic (**Figure A.2**) methods using two independent raters (Niemann et al., 2024b), with Bland-Altman analysis demonstrating high agreement between raters showing no bias and low limits of agreement. Because of the high agreement, we randomly choose one rater for the overall method comparison and provide data of the second rater in the Appendix.

### 2.5 Method comparisons

To evaluate the difference or agreement between the three-electrode center coordinate extraction methods (fully automated, semi-automated, manual), we computed the Euclidean Norm of the *x*/*y*/*z* coordinates (formula: √(*x*²+*y*²+z²)) for each subject, session, run, and electrode.

#### 2.5.1 Difference between methods

The ground truth data set consisted of 54 expert-segmented images. For comparative analysis, all other methods (fully automated, semi-automated, manual) were restricted to matching subject/session/run combinations present in this ground truth set. Coordinate extraction followed the same protocol (i.e., using center of mass calculations). We analyzed the Euclidean distances between electrode localizations using linear mixed-effects models (for details see **Table 1**) implemented in R (version 4.4.3; R Core Team, 2023) with the lme4 package (Bates et al., 2015). Model selection was conducted by comparing information criteria across candidate specifications. The final model demonstrated superior fit with an AIC (Aikake Information Criterion) of 3679.7 and BIC (Bayesian Information Criterion) of 3716.3, with substantially smaller AIC than simpler alternatives (ΔAIC > 17). The selected model structure incorporated three nested random effects to properly account for the hierarchical nature of the data: area-level variability (variance = 26.08), subject-level variability within areas (variance = 4.50), and electrode-level variability within subjects (variance = 66.68). This hierarchical structure reflected our experimental design, where electrodes were nested within subjects, and subjects were nested within anatomical areas. This analytical approach provided robust estimates of method differences, while properly accounting for the nested data structure inherent in our experimental design. The model was fitted using restricted maximum likelihood estimation with the bobyqa optimizer (Powell, 2009) to ensure convergence. Fixed effects for the factor “extraction method” were evaluated using Satterthwaite’s approximation for degrees of freedom (Satterthwaite, 1946). Post hoc comparisons were conducted using estimated marginal means (Lenth, 2023) with Tukey adjustment for multiple comparisons (see **Table 2** for results). All analyses used Kenward-Roger degrees of freedom approximation and a 95% confidence level.

**Table 1:** Linear Mixed Model Results. Linear mixed-effects model results for Euclidean norm distances across segmentation methods. Fixed effects: Intercept (ground truth reference) and method coefficients. Random effects: Nested variance structure for Area, Subject, and Electrode. Confidence intervals (CI) at 95% level. SE (standard error)

| Model: Euclidean_Norm ~ Method + (1 \| Area) + (1 \| Area:Subject) + (1 \| Area:Subject:Electrode) |  |  |  |  |  |
| --- | --- | --- | --- | --- | --- |
| Term | Estimate | SE | CI (low) | CI (high) | p-value |
| (Intercept) | 93.440 | 3.068 | 80.170 | 106.710 | 0.001 |
| Method: fully-automated | 0.206 | 0.222 | -0.230 | 0.642 | 0.35 |
| Method: semi-automated | -1.123 | 0.215 | -1.545 | -0.701 | <0.001 |
| Method: manual | 6.371 | 0.215 | 5.949 | 6.792 | <0.001 |

**Table 2:** Pairwise Comparisons Between Methods. Pairwise comparisons of Euclidean norm distances between segmentation methods using estimated marginal means (EMMs). Contrasts were evaluated via t-tests with Tukey adjustment for multiple comparisons. The ground truth method served as reference level. Significant differences (p < .001) were observed between manual and all other methods. SE (standard error)

| Contrast of EMMs | Estimate | SE | t-ratio | p-value |
| --- | --- | --- | --- | --- |
| ground truth vs. (fully automated) | -0.21 | 0.61 | -0.35 | 0.985 |
| ground truth vs. (semi-automated) | 1.13 | 0.59 | 1.91 | 0.223 |
| ground truth vs. manual | -6.37 | 0.59 | -10.80 | <.001 |
| (fully automated) vs. (semi-automated) | 1.34 | 0.61 | 2.19 | 0.127 |
| (fully automated) vs. manual | -6.15 | 0.61 | -10.05 | <.001 |
| (semi-automated) vs. manual | -7.49 | 0.59 | -12.67 | <.001 |

#### 2.5.2 Agreement between methods

Data from 290 images were used for pairwise comparisons via Bland-Altman analysis and intraclass correlation coefficients (ICC). These images were originally collected for network performance testing and included 340 images, of which 4 electrodes were correctly segmented and labeled in 323 images. The remaining 33 images were not correctly labeled.

##### 2.5.2.1 Bland-Altman analysis

Bland-Altman plots (Bland and Altman, 1999) visualize the agreement between two quantitative measurements by plotting their differences against their averages, highlighting bias and limits of agreement. Non-parametric Bland-Altman plots (**Figure 2**) were employed after Shapiro-Wilk and Kolmogorov-Smirnov tests confirmed non-normal distributions of differences between methods (*p* < 0.05 for all pairwise comparisons, **Table E.1**; Giavarina, 2015) . The x-axis represents the average of two methods, while the y-axis represents difference between them. The median difference or bias is calculated by the sum of all differences divided by the number of pairs (illustrated in **Figure 2** as dashed blue line). This value indicates whether one method consistently over- or underestimates electrode coordinates compared to the other. Limits of Agreement (LoA) are shown as dashed red line and were calculated as follows: LoA=median±1.5*IQR (Interquartile Range).

**Figure 2:**
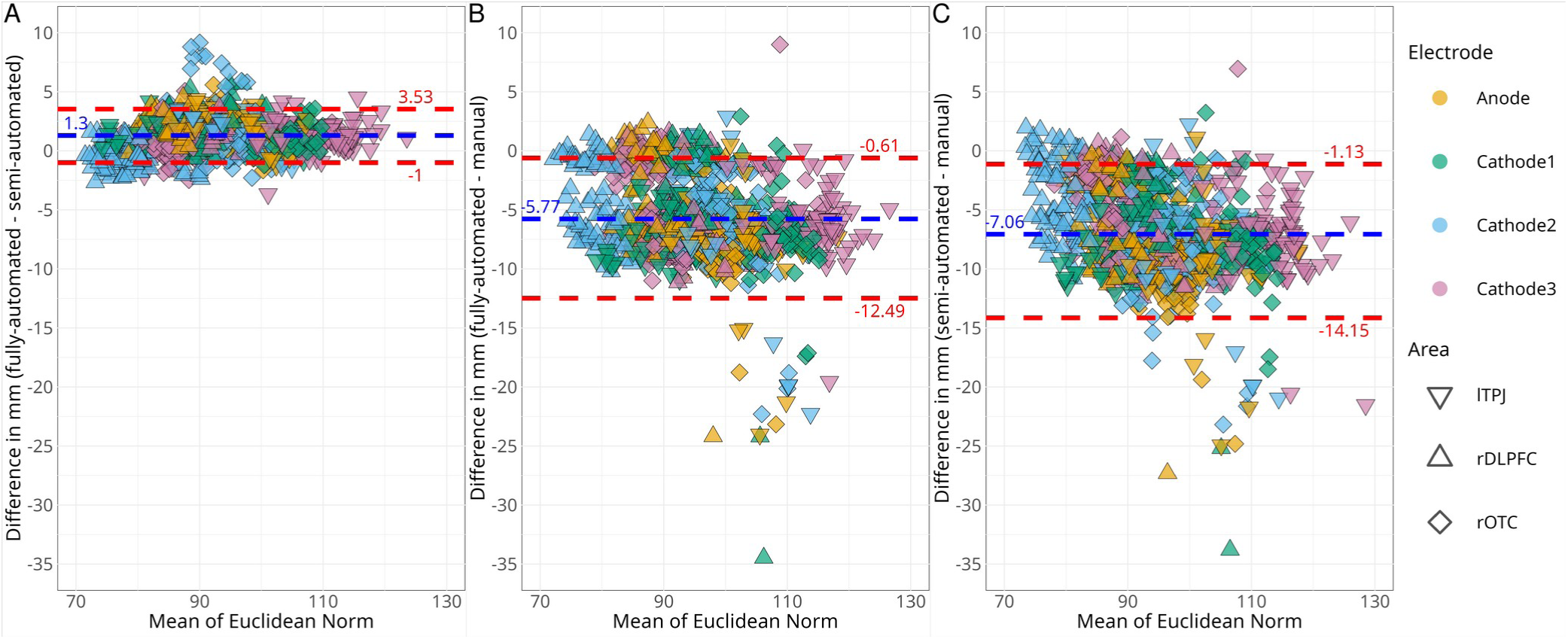
Pairwise method comparison using Bland-Altman plots. Non-parametric Bland-Altman plots comparing electrode coordinate extraction methods across three montage areas. Differences (y-axis) between paired methods are plotted against their mean Euclidean distance (x-axis) for each electrode. A) Fully vs. semi-automated extraction. B) Fully automated vs. manual extraction. C) Semi-automated vs. manual extraction. Blue dashed line: median difference (bias); red dashed lines: limits of agreement (LoA). lTPJ (left temporoparietal junction), rDLPFC (right dorsolateral prefrontal cortex), rOTC (right orbito-temporal cortex)

##### 2.5.2.2 Intraclass correlation coefficients (ICC)

To statistically evaluate the agreement between the three electrode coordinate extraction methods, we calculated pairwise ICCs using a two-way mixed-effects model for absolute agreement (see **Table 3**). Because there was (1) minimal within-subject variability across nested sessions and runs (Niemann et al., 2024b) and (2) minimal between-subject variability at the cluster level (electrode data: unpublished; area data: (Niemann et al., 2024b), where cluster-level variance effectively represents aggregated subject-level effects (Lüdtke et al., 2011; Moerbeek, 2004), we averaged individual electrode coordinates (i.e. anode or cathodes) per subject (over all sessions and runs). This approach separately accounts for both nested (session/run) and cluster-level (subject) variance components, preventing estimation bias in unbalanced designs (Moerbeek, 2004). This yielded a single representative coordinate set per subject and method. For each pairwise comparison (fully automated vs. semi-automated, fully automated vs. manual, semi-automated vs. manual), we applied the ICC(2,k) model (*two-way random-effects for agreement, mean of k raters*) via the irr package (Gamer et al., 2019) in R. This model accounts for subject variability (random effect) and systematic bias between methods (fixed effect). ICC values were interpreted as follows: <0.4 (Poor), 0.4-0.59 (Fair), 0.6-0.74 (Good), and ≥0.75 (Excellent) agreement (Koo and Li, 2016; Liljequist et al., 2019) . Confidence intervals (95%) and *p*-values were reported to quantify uncertainty. To enable direct statistical comparison of ICC values across method pairs, we applied Fisher’s z-transformation to all ICC estimates (see **Table 4**; Fisher, Ronald Aylmer, 1921)). P-values were adjusted for multiple comparisons using the Holm method (Holm, 1979).

**Table 3:** ICC Values with 95% Confidence Intervals. Pairwise ICC comparison between all 3 electrode coordinate extraction methods (full, semi-automated and manual). Results of ICCs are shown for all electrodes and each electrode individually (Anode, Cathode_1-3_). ICC (Intraclass Correlation Coefficient)

| Electrode | fully-automated vs semi-automated | fully-automated vs manual | semi-automated vs manual |
| --- | --- | --- | --- |
| All | 0.990 (0.924-0.996) | 0.901 (-0.102-0.976) | 0.862 (-0.096-0.966) |
| Anode | 0.967 (0.394-0.991) | 0.800 (-0.163-0.949) | 0.725 (-0.101-0.929) |
| Cathode1 | 0.989 (0.805-0.997) | 0.874 (-0.117-0.970) | 0.836 (-0.071-0.962) |
| Cathode2 | 0.989 (0.974-0.994) | 0.906 (-0.100-0.978) | 0.873 (-0.124-0.970) |
| Cathode3 | 0.997 (0.932-0.999) | 0.942 (-0.052-0.987) | 0.920 (-0.066-0.983) |

Note: ICC Interpretation: <0.4 Poor, 0.4-0.59 Fair, 0.6-0.74 Good, ≥0.75 Excellent (green)

**Table 4:** ICC Comparisons with Holm-Adjusted p-values. Statistical comparison of ICC values using Fisher’s z-transformation. Table shows p-values for pairwise reliability comparisons between extraction methods across electrode groups. Significance levels: ***p<0.001, **p<0.01, *p<0.05.

| Comparison | All | Anode | Cathode<sub>1</sub> | Cathode<sub>2</sub> | Cathode<sub>3</sub> |
| --- | --- | --- | --- | --- | --- |
| ICC (fully automated, semi-automated) vs ICC (fully-automated, manual) | 1.0e-08*** | 3.6e-06*** | 6.1e-10*** | 9.3e-08*** | 1.2e-12*** |
| ICC (fully automated, semi-automated) vs ICC (semi-automated, manual) | 5.2e-11*** | 3.8e-08*** | 6.3e-12*** | 1.2e-09*** | 2.8e-15*** |
| ICC (fully automated, manual) vs ICC (semi-automated, manual) | 0.38 | 0.36 | 0.46 | 0.43 | 0.40 |

Note: Significance codes: \*\*\* $p < 0.001$ , \*\* $p < 0.01$ , \* $p < 0.05$ $p$ -values adjusted using Holm's method

## 3 Results

### 3.1 Network metrics

The network achieved a DSC of 0.76, HD = 36.76. With post-processing of segmented images 95% of images contained exactly four electrodes, i.e., 323 out of 340 images were correctly segmented (see also **Figure 2C3** for segmentation examples and **Figure 3** for extracted coordinate examples). This was the case for 323 individual images, while at least one electrode was omitted in 12 images (3.71%). In two images no electrodes were identified (0.26%). In addition, automatic labeling of identified electrodes as center anode or cathodes_1,2,3_ (for details see below) was performed, to facilitate extraction method comparisons. In 323 of the correctly segmented 323 images, all four electrodes electrodes were correctly labeled. Those images comprised a total of 1.324 electrode positions from overlapping participants and imaging sessions that were used for direct comparisons of electrode position extraction accuracy achieved with the three respective approaches.

**Figure 3:**
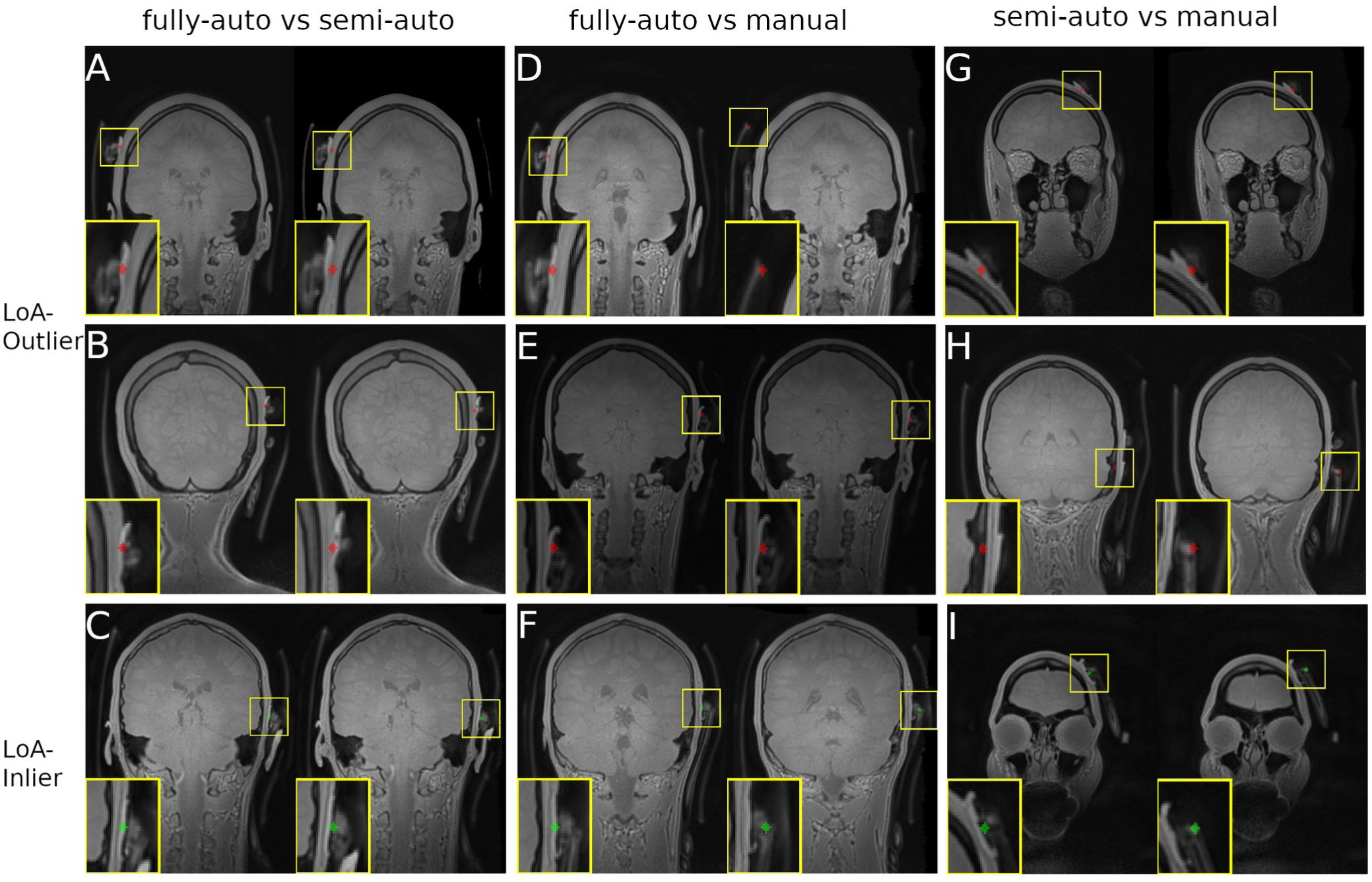
Representative image examples of the Bland-Altman plots. Representative examples of electrode coordinate classification from Figure 2. Top/Middle rows: Outliers (red spheres) outside Limits of Agreement (LoA). Bottom row: Inliers (green spheres) within LoA.

### 3.2 Method comparison

#### 3.2.1 Difference between methods

##### 3.2.1.1 Linear Mixed Model

Results in **Table 1** revealed significant differences between localization methods of 52 images. The manual method showed substantially larger mean deviations from the ground truth (up to 6 mm; β = 6.37 mm, SE = 0.22, p < .001) compared to other approaches. The semi-automated method exhibited smaller but still significant mean differences of ∼1 mm (β = -1.12, SE = 0.22, p < .001), while the fully automated method did not significantly differ from the ground truth (p = 0.35).

##### 3.2.1.2 Estimated Marginal Means (EMM)

Analyses of EMM in **Table 2** of the linear mixed model in **Table 1** revealed significant differences between methods. Pairwise comparisons with Tukey adjustment showed that the manual method (M = 99.8, 95% CI: [86.6, 113]) differed significantly from all others (all p < .0001, with t-values ranging from 27.66 to 34.75). The semi-automated method (M = 92.3, 95% CI: [79.1, 105]) yielded significantly lower estimates than both ground truth (M = 93.4, 95% CI: [80.3, 107]; t(589) = 5.23, p < .0001) and the fully automated method (M = 93.6, 95% CI: [80.5, 107]; t(589) = 5.97, p < .0001). No significant difference was found between ground truth and the fully automated method (t(589) = -0.93, p = .789).

#### 3.2.2 Agreement between methods

##### 3.2.2.1 Bland-Altman plots

Non-parametric Bland-Altman plots of 1160 data points (290 images *4 electrodes (Anode, Cathode_1-3_)). For each pairwise method comparison across all montages and electrodes are shown in **Figure 2**. The comparison between fully and semi -automated methods revealed the smallest bias and narrowest LoAs (bias = 1.30 mm; LoA: [−1.00, 3.53]), with 90.69% of data points (1052/1160) falling within the LoA (**Figure 2A**). In contrast, comparisons involving manual extraction exhibited larger biases and greater variability. The fully automated vs. manual comparison showed a bias of −5.77 mm (LoA: [−12.49, −0.61]; **Figure 2B**), while the semi-automated vs. manual comparison had a bias of −7.06 mm (LoA: [−14.15, −1.13]; **Figure 2C**). Only 87.33% and 91.47% of points, respectively, fell within the LoA for these comparisons.

##### 3.2.2.2 Visual inspection of extraction-method performance

Visual inspection of images outside the limits of agreement (LoAs) in **Figure 2** (all outliers available https://nextcloud.uni-greifswald.de/index.php/s/HmtA4wtqkbkaqy7, accessed July 29.2025) revealed distinct error patterns that are described below.

###### 3.2.2.2.1 Fully- vs. Semi-Automated Extraction (Figure 3A–C, Figure 2A)

Among 108 outliers (i.e. individual electrodes; out of 1160 data points (Figure 2)), 24 showed minimal deviations of the extracted electrode centers during visual inspection of extracted electrodes for both methods (**Figure 3A,B**). In another eight cases, the fully automated extraction placed electrodes slightly off-center, whereas semi-automated extraction performed correctly (**Figure 3B**). The remaining 72 outliers were explained by electrode center deviations during the semi-automatic extraction (**Figure 3A**). An example of method differences within the Limits of Agreement (LoA) is also shown to illustrate minor deviations from the actual position (**Figure 3C**).

###### 3.2.2.2.2 Fully-Automated vs. Manual Extraction (Figure 3D–F, Figure 2B)

Of 147 outliers, 20 cases exhibited drastic misplacement that occurred during manual extraction (e.g., a −34.5 mm difference, **Figure 3D**). Four cases showed minor errors with the fully automated extraction method (**Figure 3E**), while 101 cases revealed similar center coordinates with minimal error for visual inspection for both methods (**Figure 3E**, difference = 1.2 mm). The remaining 22 cases showed deviations exclusively for the manual extraction method, ranging from mild to severe (**Figure 3G and 3H**), while the fully automated extraction accurately identified the electrode center. Notably, many differences like in **Figure 3E** were classified as outliers due to manual extraction bias (−6.55 mm), shifting the LoA away from near-zero differences. Manual extraction frequently placed coordinates on the electrodes plug (For visual clarification of the bias see **Figure 3F, right**), which - despite being incorrect - appeared as correct placement (**Figure 3E, right**).

###### 3.2.2.2.3 Semi-Automated vs. Manual Extraction (Figure 3G–I, Figure 2C)

Among 99 outliers, 20 cases exhibited drastic misplacement during manual extraction (e.g., a −34.5 mm difference, **Figure 3D**) and manual extraction errors dominated overall. In 67 cases, both methods deviated similarly, with the manual bias primarily accounting for outlier classification (**Figure 3G**). Only two outliers stemmed from inaccuracies during semi-automated method showing misplacement in the opposite direction of the manual extraction (**Figure 3H, left**). Hence, differences in the Bland-Altman plot were more pronounced relative to the comparison of fully automatic vs. manual extraction (**Figure 2C**). Note, semi-automated coordinates that fall outside the LoA compared to the fully automated method still remain within the LoA when benchmarked against manual annotations. Since manual error introduces more pronounced misplacement, the semi-automated method demonstrates relatively better performance.

###### 3.2.2.2.4 Key Error Patterns

The most pronounced errors between fully and semi-automated extraction occurred when fully automated segmentation correctly identified electrodes, while semi-automated coordinates erroneously positioned them into the brain (**Figure 3A,H**), creating a small positive bias of 1.3 mm (**Figure 2A**). Manual extraction frequently misallocated coordinates onto electrode plugs **(Figure 3F,I)** or on the inflatable air cushions (**Figure 3D**), contributing to substantial outliers in **Figures 2B and 2C**.

##### 3.2.2.3 ICC

The inter-method reliability analysis using ICCs revealed distinct patterns across electrode types and measurement approaches (see **Table 3**). Fully- vs semi-automated methods had excellent agreement across all electrodes (i.e., anode and cathodes; ICC=0.990, 95% CI [0.924-0.996]). Comparisons of fully automated vs manual extractions revealed an excellent point estimate (ICC=0.901), but also wide confidence intervals (95% CI: [-0.102-0.976]). Negative lower bounds suggest potential reliability concerns. Comparison of semi-automated vs manual extractions showed good agreement (ICC=0.862), but also wide 95% CI: [-0.096-0.966]. Fully-automated vs semi-automated comparisons consistently achieved excellent reliability across all electrodes (ICC≥0.967), with superior extraction for the fully automated network. In contrast, negative lower bounds in comparisons involving manual extractions suggest potential model/data limitations and comparisons with manual extraction of the anodes showed the poorest agreement (ICC=0.800-0.725).

##### 3.2.2.4 Statistical Comparison of ICC Values Using Fisher’s z-Transformation

Fisher’s z-transformation was applied to compare ICCs between method pairs across electrode groups (**Table 4**). The ICCs for fully vs. semi-automated extractions were significantly higher (*p* < 0.001) than those for fully automated vs. manual and semi-automated vs. manual across all electrodes (All: *p* < 0.001; Anode: *p* < 0.001; Cathodes 1–3: *p* < 0.001). This indicates superior agreement between the two (semi)automated methods compared to comparisons with manual extraction. Differences between fully automated or semi-automated with manual ICCs were comparable for all electrodes (all: *p* = 0.38; anode: *p* = 0.36; cathodes 1–3: *p* = 0.40–0.46). This suggests that while both automated methods disagreed with manual extraction to a similar degree, their agreement with each other remained strong. Consistent with **Table 3**, the Fisher-transformed comparisons highlight that (1) fully vs. semi-automated ICCs consistently reached "excellent" agreement (ICC ≥ 0.75) for all electrodes and (2) automated vs. manual ICCs varied more widely, with wider confidence intervals (e.g., fully-automated vs. manual: −0.102 to 0.976), due to errors during manual extraction.

## 4 Discussion

This study introduced a fully automated deep learning approach, trained on highly accurate, manually segmented electrodes serving as ground truth, to extract tDCS electrode positions from structural MRI data. Its performance was compared to manual and semi-automated approaches using linear mixed-effects models, Bland-Altman analysis, and ICC reliability testing. Statistical comparison of methods revealed significant mean deviations from the ground truth for manual and semi-automated electrode coordinate extraction, whereas the fully automated method showed no significant deviation. Overall, results suggested that fully and semiautomated methods exhibit superior consistency and accuracy compared to manual extraction, though challenges persist in specific scenarios. Overall, segmentation performance of the novel deep learning approach was excellent and in 95% of the input images all electrodes were correctly identified, while this approach offered a high degree of efficiency. Our specific approach and possible adaptations accounting for different tDCS set-ups, have potential to enhance efficiency and accuracy of electrode extraction in focal tDCS-fMRI studies. From a practical point of view, electrode extraction via deep learning approaches can facilitate inspection of targeting accuracy and the outcomes can be used to improve the validity of individualized current modeling studies, which is particularly relevant for tDCS-fMRI studies investigating dose-response relationships.

Importantly, while electrode positioning errors can change the overall electric field distribution (e.g.,Indahlastari et al., 2023), those may not substantially affect current dose in the immediate target regions for conventional tDCS montages (Grignard et al., 2023; Niemann et al., 2024a). Thus, validation of electrode placement accuracy by visual inspection may be sufficient in many scenarios (Antonenko et al., 2019; Darkow et al., 2017; Meinzer et al., 2012), even though segmentation of electrodes from MRI/CT images is recommended (Indahlastari et al., 2021; Opitz et al., 2018). For focal set-ups, electrode positioning validation and use of actual electrode positions in computational models is more critical, because minor deviations from intended positions can lead to significant dose reductions in the target regions (Niemann et al., 2024a). Notably, even though the majority of focal tES-fMRI studies used imprecise scalp-based methods for the initial positioning of electrodes (e.g., 10-20 EEG system), placement accuracy was frequently not validated (Hampstead et al., 2020; Leaver et al., 2023; Müller et al., 2023; Rauh et al., 2023). Moreover, even if the initial electrode positioning is optimized (e.g., by neuronavigation; Jog et al., 2021; Niemann et al., 2024a), intrascanner tDCS studies are at high risk for electrode displacement during subsequent positioning of participants in the scanner or participant movement inside of the bore (see Meinzer et al., 2024). Hence, verification of electrode positions before (to confirm placement accuracy inside the scanner) and after functional imaging (to investigate potential electrode drift) is recommended for tES-fMRI studies. Given the increased interest in studying the neural mechanisms by which (focal) tDCS modulates human brain functions with fMRI and individualized current modeling in larger cohorts (e.g., www.memoslap.de), development of novel automated methods for electrode position extraction and their validation are urgently needed to ensure efficient and precise workflows.

So far, however, only two focal tDCS-fMRI studies have used either manual (Niemann et al., 2024a) or semi-automatic (Niemann et al., 2024b) approaches to extract electrode positions from structural MRI data, which is time consuming and results can be affected by human error. The latter was confirmed in the present study by the statistical comparison of methods that revealed the mean deviations from ground truth for the manual approach (up to 6 mm). In contrast, semi-automated methods showed smaller but still significant differences (∼1 mm), whereas the fully automated network approach performed comparably to ground truth. Analysis of estimated marginal means (**Table 2)** further supported these findings, with pairwise comparisons confirming that the manual method differed significantly from all others. Notably, no meaningful difference was observed between network-based extraction and ground truth, emphasizing the validity of our approach. Nonetheless, since most training and validation data were derived from the same ground truth data set, the network may have learned to fit this specific data rather than generalize to a broader distribution. However, the overall robust performance of the network (i.e., four electrodes were correctly identified in 95% of the 340 images) does not support this assumption.

While additional network features may improve edges, a DSC of 0.76 suggests that AGs already capture the most salient image patterns. In addition, DSC performance of the network was superior (Zhang et al., 2023) or in the same range compared to other fMRI-based small structure segmentation tasks (Fassia et al., 2024; Lim et al., 2021; Vaidyanathan et al., 2021), even though the moderate Hausdorff distance suggests occasional outliers in boundary precision (for visualization see **Figure 3E left**). Future improvements should therefore prioritize edge-case refinement, especially for electrodes near anatomical boundaries or imaging artifacts. Indeed, visual inspection revealed that most false positives occurred at electrode boundary structures, while some unsegmented electrodes exhibited significant deformation or unusual curvature, potentially due to study specific methods used for head immobilization or to improve tDCS impedance. Incorporating additional feature maps in the network architecture (for visualization see **Figure B.1)** or feature map attention (Sang et al., 2023) may further improve edge detection accuracy and minimize false positives. In the present study, memory limitations precluded such optimizations.

Inspection of Bland-Altmann plots showed the highest agreement between fully vs. semi-automated extractions, with 90.79% of data points within LoA). In contrast, manual extraction introduced significant bias and variability, most pronounced for comparison with the network approach and mainly in the x dimension (Figure C.1B,C; for further details see also Appendix C, Figure C.1-4). Notably, 15–20% of manual comparisons fell outside LoAs, with visual inspection revealing systematic errors (e.g., center coordinates placed on electrode knobs or air cushions, Figure 3D,F), while semi-automated errors typically involved off-center deviations (Figure 3A,H). A similar pattern was observed for the second rater, with Bland-Altmann plots (Figures D.1) demonstrating comparable trends in agreement and systematic errors. These findings suggest that extractions involving manual steps are prone to systematic errors. Finally, the ICC analysis confirmed excellent reliability (≥ 0.97) between semi-and fully automated methods, and superiority compared to manual extraction. (Semi- and fully) automated vs. manual ICCs showed wider confidence intervals, due to higher variability of manually extracted coordinates. The anode exhibited the poorest manual agreement for automated vs. manual ICCs, which could be due to the fact, that the anodes were most consistently covered by the inflatable cushion. Crucially, the degree of deviation between both (semi- and fully) automated methods from manual extraction was comparable, suggesting that manual variability, rather than algorithmic flaws, drove discrepancies. Therefore, automated methods reduce human error and significantly decrease bias, particularly in the *x* dimension for the given montages, where manual placement showed systematic deviations.

Another key advantage of the novel network approach is the elimination of preprocessing steps required by alternative methods, such as 2D-to-3D transformation in semi-automated pipelines or air/threshold adjustments in manual segmentation. In addition, our framework offers a scalable solution for verifying electrode placement in tDCS-fMRI studies, ensuring computational models of current flow are based on actual (rather than assumed or falsely identified) electrode positions. Future adaptations could accommodate different multi-array or ring tDCS setups (Dmochowski et al., 2011; Gbadeyan et al., 2016; Khan et al., 2022) or integration with real-time quality control pipelines (Dupuis et al., 2024; Specktor-Fadida et al., 2025). By improving the accuracy and efficiency of electrode extraction, this approach may be suited to enhance the validity of tDCS-fMRI studies, particularly those probing dose-response relationships or individual variability in stimulation effects. The narrow LoAs and high ICCs between automated approaches support their use in large-scale tDCS-fMRI studies to minimize human error and improve reproducibility. Furthermore, the negative lower bounds ICC confidence intervals and extreme outliers for the manual extraction method question its reliability as a standalone reference. Protocols may benefit from hybrid verification (e.g., automated extraction with manual oversight for ambiguous cases). Semi-automated deviations from electrode centers (**Figure 3A**) and manual biases toward the electrode plug (**Figure 3D**) highlight specific failure modes for algorithm improvement.

In sum, both fully and semi-automatic approaches outperformed manual extraction, by showing superior alignment with actual electrode positions (ground truth). Our network showed high precision in electrode extraction in a wider data set, when it was able to detect them. Especially for larger scale studies that require extensive data extraction, a network-based approach is preferable to improve efficiency.

## 5 Limitations

Our study is limited by the relatively small ground truth data set (54 images), which was sufficient for training the network. Generalization performance (95%; 323 out of 340 correctly identified 4 electrodes, see section 3.1) may be further improved by edge-case refinement to address occasional boundary inaccuracies, particularly near anatomical artifacts. Moreover, network generalization can be improved with minimal additional data from other montages, suggesting that sufficient training across diverse data sets and montages may allow generalization to most variations. We tested this approach using our 54 highly accurate, manually segmented images and augmented the data set with 18 additional highly accurate segmentations from five independent projects of the research unit with different montages and target regions (∼3 new segmented images per project). Out of a total of 1,823 images (72 manually segmented images or training data are already excluded) 91.39% (1,666 segmented images) correctly identified four electrodes. These results demonstrate that our network generalizes beyond the initial training data and can adapt to new montages with only a small amount of additional annotated data. However, memory constraints prevented implementation of expanded feature spaces that could further improve edge detection. While the proposed attention-gated U-Net demonstrates robust performance in segmenting small electrodes on PETRA MRI scans, its efficacy on conventional T1-weighted images remains limited (data not shown). This discrepancy arises from the poorer contrast-to-noise ratio between electrodes and surrounding tissues in conventional T1 scans, where the electrode gel intensity often matches the signal of the head, while the electrodes themselves blend into the background. The T1 images also show more tissue structures inside the head, making them more demanding target images. Consequently, the network fails to learn discriminative features, resulting in negligible segmentation improvement. Hence, we currently recommend to acquire PETRA scans in tDCS-fMRI, to facilitate potential implementation of a network-based electrode extractions.

## 6 Conclusion and Outlook

Although electrode placement verification is recommended in current tDCS-fMRI reporting guidelines (Ekhtiari et al., 2022), this has rarely been implemented. Current validation checks in tDCS-fMRI studies predominantly involved visual inspection and seldom manual electrode segmentation, manual electrode extraction or neuronavigated validation, because these methods are time-consuming, prone to human error, or lack standardization. We therefore developed and validated an open-source network for automated electrode segmentation and coordinate extraction which should be adopted in all future studies of focal tDCS-fMRI.

In addition to being accurate and efficient, another key advantage of this approach is that the network can be retrained with limited additional data to improve its generalizability. When the network fails to segment a new montage, only few precisely segmented ground truth data sets are needed for retraining (Tizpaz-Niari and Kreinovich, 2023) to improve generalization. While additional network features may improve edges, a DSC of 0.76 suggests that AGs already capture the most salient image patterns and additional marginal gains may not justify memory costs unless boundary precision is critical. Therefore, targeted feature augmentation could be implemented. Post-Processing could be advanced by applying a lightweight edge-refinement CNN after AG-based segmentation (Takikawa et al., 2019). Optimizing AGs while testing if AGs benefit more from feature diversity (e.g., texture/edge channels) vs. feature quantity. To address the challenges of electrode segmentation in T1-weighted MRI, future work could employ intensity normalization or synthetic contrast enhancement (Santini et al., 2018), e.g., via generative adversarial networks or diffusion models to improve electrode visibility. Such approaches may compensate for the inherently low contrast-to-noise ratio (CNR) between electrodes and background tissues in conventional T1 scans, enabling more reliable automated segmentation.

## Acknowledgments

We thank Hanka Schalinksi for her diligent work in manually segmenting 40 intracranial electrodes, which contributed significantly to this study. We also thank Sophie Dabelstein and Kira Hering for help with data extraction. The authors acknowledge the use of AI (google gemini: 2.5 flash) for optimizing the wording and grammar of parts of this manuscript, but it was not used to generate any of the core scientific content.

## Ethics Statement

The study was approved on February 20, 2020, by the local ethics committee (reference number BB 172/19) and was conducted in accordance with the Helsinki Declaration. Written informed consent was obtained prior to inclusion in the study.

## Data Statement

Raw structural MRI data, including Pointwise Encoding Time Reduction with Radial Acquisition (PETRA) scans, were used for this study. Due to their nature as human subject data, these raw images will not be shared publicly to protect participant privacy.

All code used for training the deep learning network (a 3D Attention U-Net) to automatically segment and extract electrode coordinates is available in a publicly accessible GitHub repository. This includes scripts for network implementation, training protocols, model evaluation, and post-processing steps. The shared code allows for full replication of the image analysis pipeline, ensuring the reproducibility of our findings if similar datasets are available.

