## Appendix for "Deep learning to overcome human error and bias in electrode position extraction in tDCS-fMRI studies"

A. Rater reliability

Rater reliability from 2 raters of both electrode extraction methods, manual (**Figure A.1**) and semi-automated (**Figure A.2**), were examined for all 3 areas for each electrode showing a small ne-glectable bias of -0.019 LoA[-2.84, 2.80] and 77.43 % of data outside the LoAs. Semi-automated however showed a slightly bigger bias of -0.14, smaller LoAs[-1.39, 1.09] and 92.65% of data in-sight the LoAs.

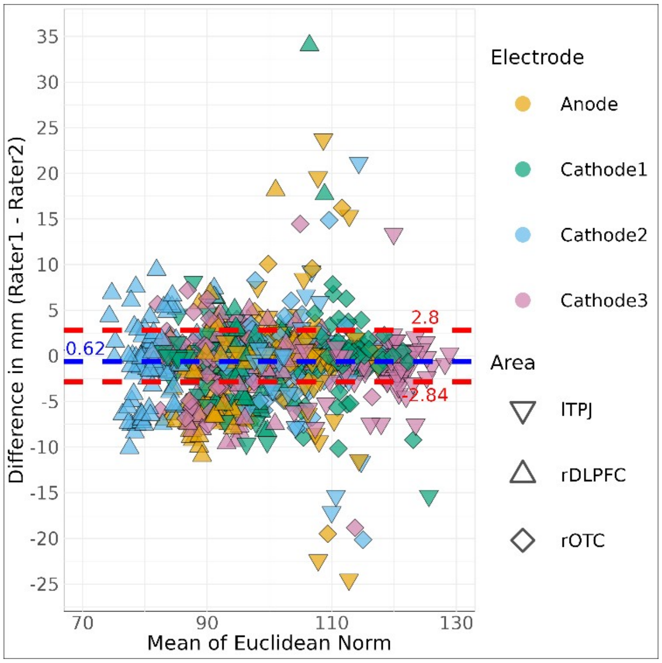


***Figure A.1**: Non-parametric Bland-Altman plots of the manual method comparing 2 raters. Non-parametric Bland-Altman plots comparing 2 raters using the manual method to extract electrode coordinates for 3 areas: lTPJ (left temporoparietal junction), rDLPFC (right dorsolateral prefrontal cortex), rOTC (right orbito-temporal cortex) and 4 electrodes of a 3x1 focal setup. Differences (y-axis) between both raters are plotted against their mean ratings (x-axis). Blue dashed line: median difference (bias); red dashed lines: limits of agreement (LoA).*

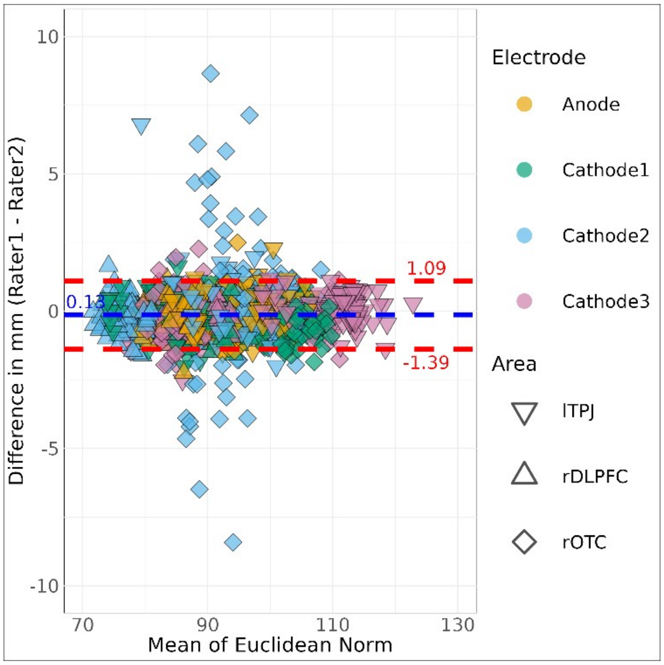


***Figure A.2****: Non-parametric Bland-Altman plots of the semi-automated method comparing 2 raters. Non-parametric Bland-Altman plots comparing 2 raters using the semi-automated method to extract electrode coordinates for 3 areas: lTPJ (left temporoparietal junction), rDLPFC (right dorsolateral prefrontal cortex), rOTC (right orbito-temporal cortex) and 4 electrodes of a 3x1 focal setup. Differences (y-axis) between both raters are plotted against their mean ratings (x-axis). Blue dashed line: median difference (bias); red dashed lines: limits of agreement (LoA).*

### B. Detailed Network Architecture

A comprehensive visualization of the network architecture is provided in **Figure B.1**, which illustrates the transformation from input images to feature maps throughout the optimization process in greater detail.

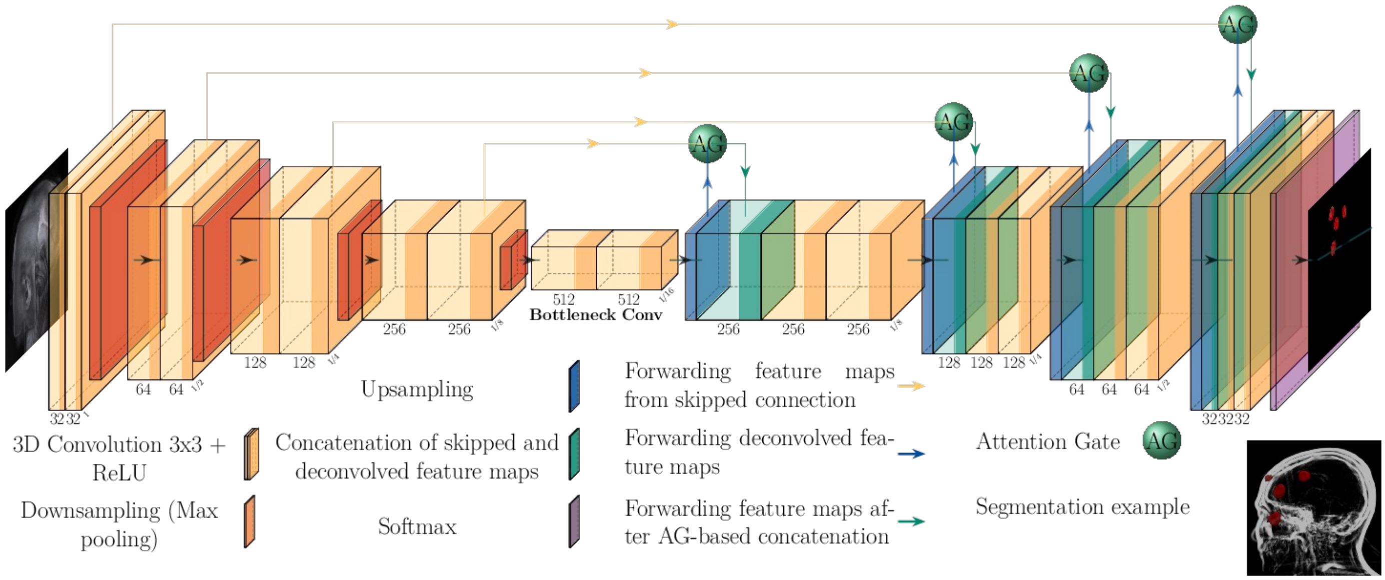


***Figure B.1***: *3D Attention U-Net architecture. 3D Attention U-Net architecture trained with Adam to minimize Dice-Focal Loss ($\lambda_{Dice}$=0.3, $\lambda_{Focal}$=0.7, γ=2.5). The network features encoder/decoder paths, skip connections with attention gates, channels (32, 64, 128, 256, 512), strides of 2, kernel size 3. ReLU (Rectified Linear Unit) was used as activation function. Achieved Dice score (DSC) = 0.76, Hausdorff distance (HD) = 36.76. (HarisIqbal88, 2018)*

3D Attention U-Net architecture. 3D Attention U-Net architecture trained with Adam to minimize Dice-Focal Loss ($\lambda_{Dice}$=0.3, $\lambda_{Focal}$=0.7, γ=2.5). The network features encoder/decoder paths, skip connections with attention gates, channels (32, 64, 128, 256, 512), strides of 2, kernel size 3. ReLU (Rectified Linear Unit) was used as activation function. Achieved Dice score (DSC) = 0.76, Hausdorff distance (HD) = 36.76. (HarisIqbal88, 2018)

### C. Dimensional Bland-Altman plots

While the Euclidean norm combines positional errors across all dimensions, the directional source of these errors becomes apparent when examining the individual coordinate axes. **Figure C.1 - 5** present separate Bland-Altman plots for the x, y, and z dimensions, revealing several important patterns in electrode placement accuracy.

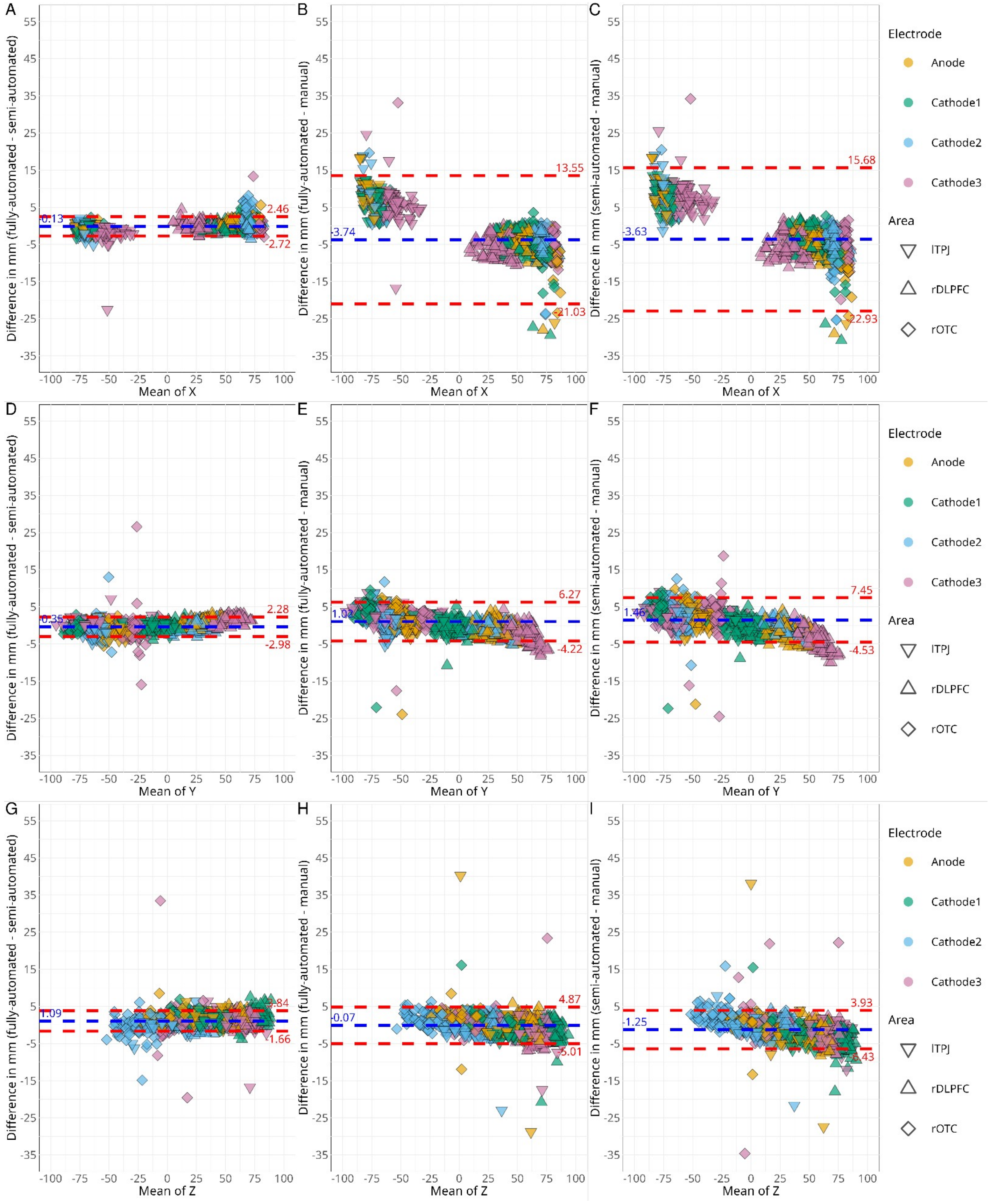


***Figure C.1**: Non-parametric Bland-Altman plots for each dimension over all montage areas. Non-parametric Bland-Altman plots comparing electrode coordinate extraction methods across three montage areas. Differences (y-axis) between paired methods are plotted against their mean distance (x-axis) for each electrode in the dimension x (**A-C**)/ y (**D-F**) and z (**G-I**). **Left column:** Fully vs. semi-automated extraction. **Middle column:** Fully automated vs. manual extraction. **Right col-***

***umn:*** *Semi-automated vs. manual extraction. Blue dashed line: median difference (bias); red dashed lines: limits of agreement (LoA).*

The analysis demonstrates that manual electrode placement introduced significant bias primarily in the x-direction, as shown in **Figure C.1B** and **C.1C**. This x-direction bias exhibited different polarities depending on the hemisphere, with positive displacement for the left temporoparietal junction (lTPJ, F**igure 3A**) and negative displacement for right-sided areas (rDLPFC and rOTC), as illustrated in F**igure 3G** and **3H**. Secondary errors in the z-direction emerged when electrodes were placed near the head's curvature, visible in **Figure 3G** and **3I**.

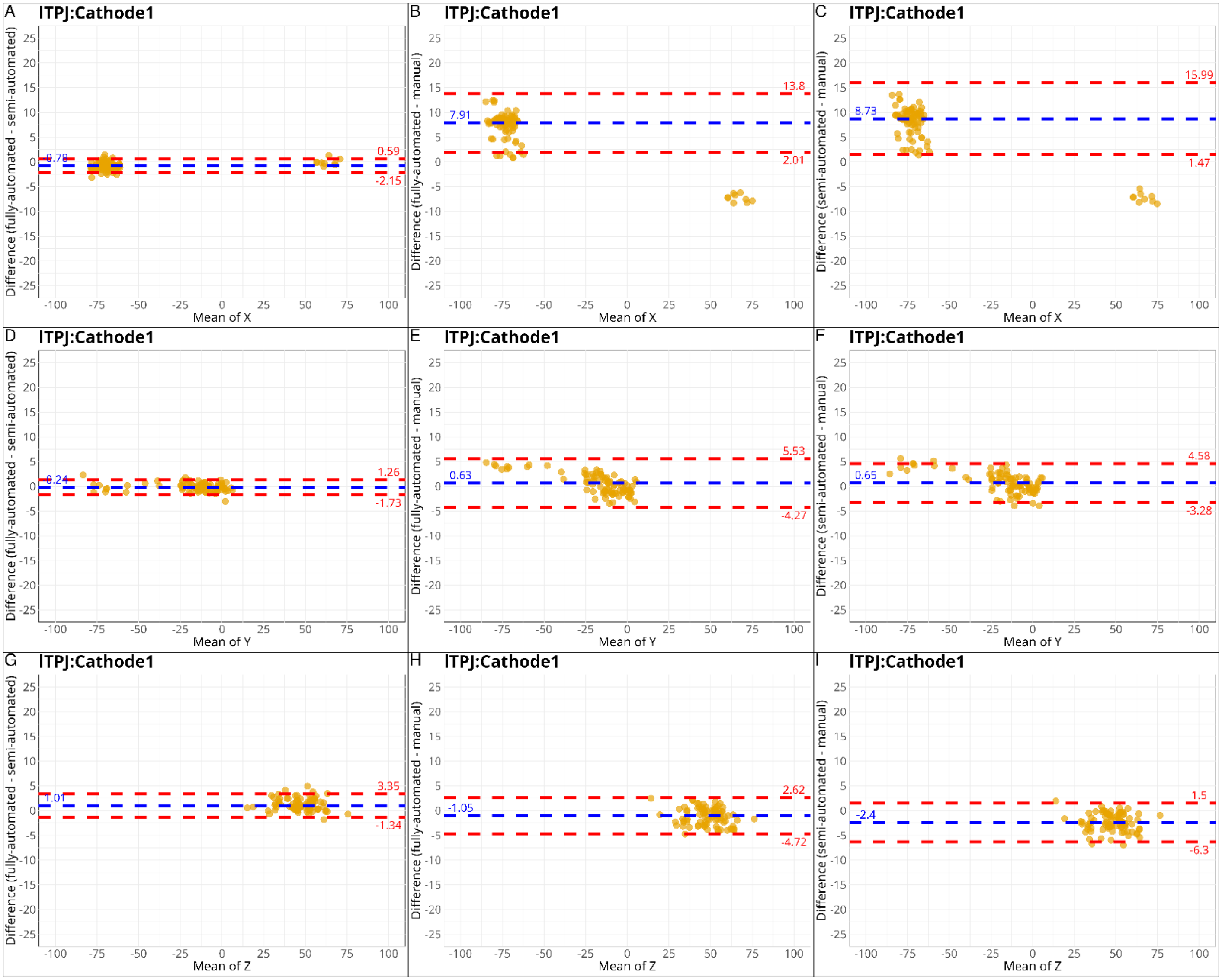


***Figure C.2****: Non-parametric Bland-Altman plots for lTPJ setup Non-parametric Bland-Altman plots comparing electrode coordinate extraction methods for montage area lTPJ (left temporoparietal junction). Differences (y-axis) between paired methods are plotted against their mean distance (x-axis) for the Cathode1 in the dimension x (****A-C****)/ y (****D-F****) and z (****G-I****).* ***Left column:*** *Fully vs. semi-automated extraction.* ***Middle column:*** *Fully automated vs. manual extraction.* ***Right column:***

*Semi-automated vs. manual extraction. Blue dashed line: median difference (bias); red dashed lines: limits of agreement (LoA).*

Comparison between methods revealed consistent advantages of automated approaches. The fully versus semi-automated method comparisons (shown in the first columns of **Figure C.2**) demonstrated substantially smaller biases and tighter Limits of Agreement compared to manual method comparisons (columns 2 and 3). Region-specific analysis showed particularly large x-direction biases in manual placement for the lTPJ, Cathode1, ranging from 7.91 mm to 8.73 mm with Limits of Agreement between 1.47 mm and 15.99 mm (**Figure C.2B** and **C.2C)**. Other dimensions in this region showed minimal errors, with maximum displacements under 2.4 mm.

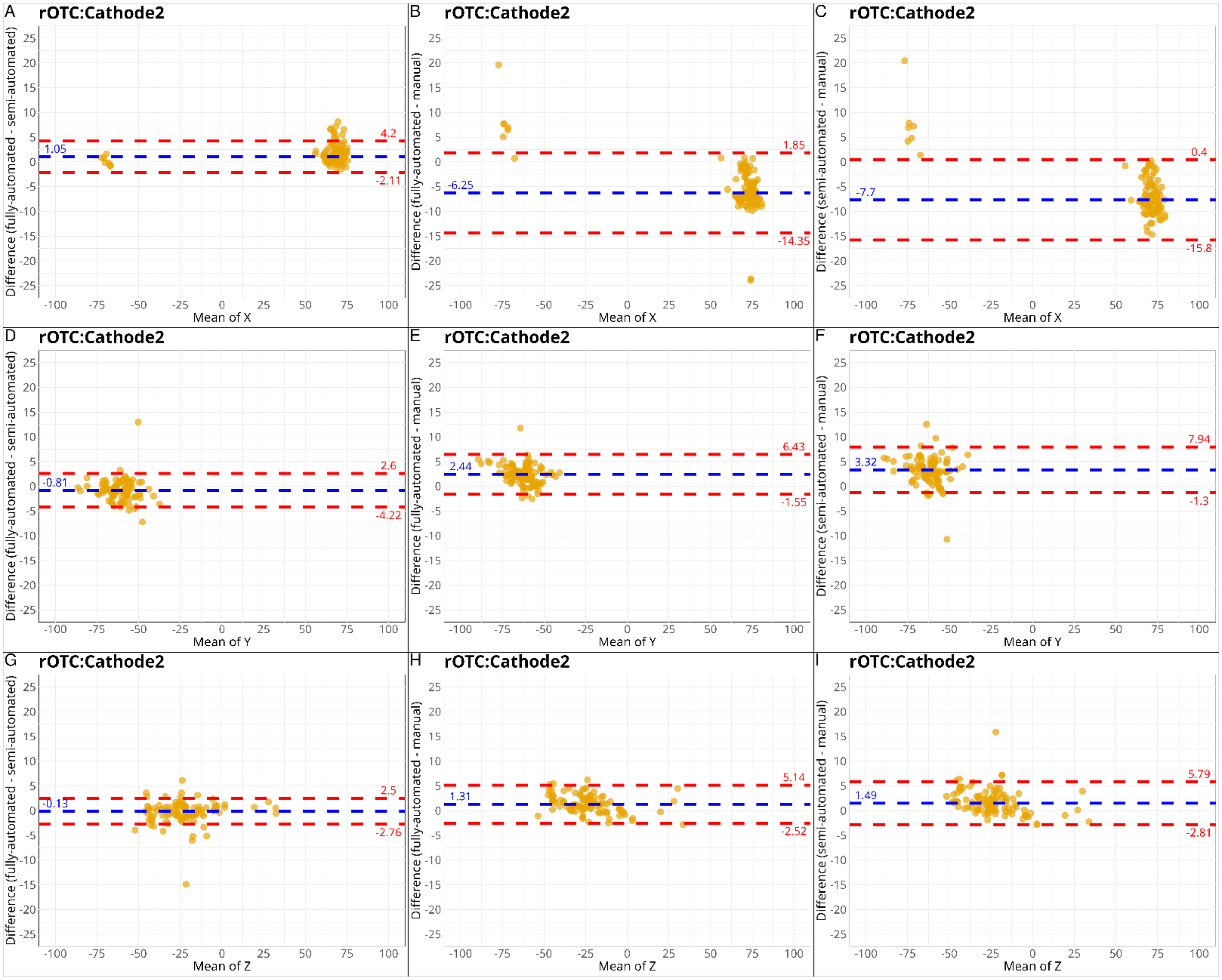


***Figure C.3**: Non-parametric Bland-Altman plots for rOTC setup. Non-parametric Bland-Altman plots comparing electrode coordinate extraction methods for montage area rOTC (right orbito-temporal cortex). Differences (y-axis) between paired methods are plotted against their mean distance (x-axis) for the Cathode2 in the dimension x (**A-C**)/ y (**D-F**) and z (**G-I**). **Left column:** Fully vs. semi-automated extraction. **Middle column:** Fully automated vs. manual extraction. **Right column:** Semi-automated vs. manual extraction. Blue dashed line: median difference (bias); red dashed lines: limits of agreement (LoA).*

For the right orbito-temporal cortex (rOTC), manual methods produced consistent negative x-direction biases of -6.25 mm to -7.7 mm, with Limits of Agreement ranging from -15.08 mm to 1.85 mm (**Figure C.3B** and **C.3C**, illustrated in **Figure 3F** and **3I**). The right dorsolateral prefrontal cortex (rDLPFC) showed more complex error patterns due to electrode orientation, with mixed x-direction biases ranging from -3.42 mm to -3.91 mm (Limits of Agreement between: -10.66 mm to 3.71 mm) and additional z-direction biases of -2.19 mm (Limits of Agreement: -8.08 mm to 3.7 mm), as documented in **Figure C.4B, B.4C**, and **B.4I**, and illustrated in **Figure 3G**.

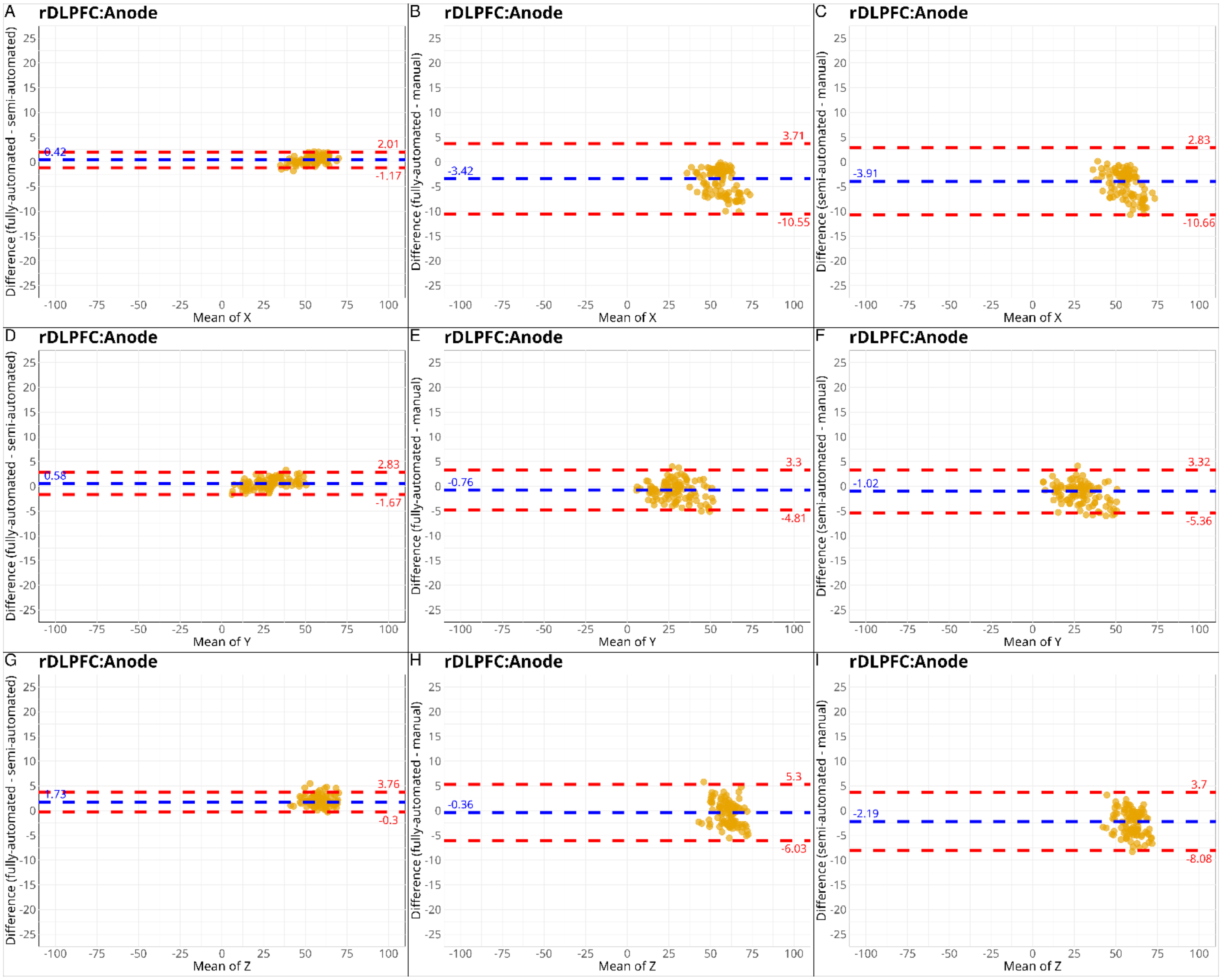


***Figure C.4**: Non-parametric Bland-Altman plots for rDLPFC setup. Non-parametric Bland-Altman plots comparing electrode coordinate extraction methods for montage area rDLPFC (right dorsolateral prefrontal cortex). Differences (y-axis) between paired methods are plotted against their mean distance (x-axis) for the Anode in the dimension x (**A-C**)/ y (**D-F**) and z (**G-I**). **Left column:** Fully vs. semi-automated extraction. **Middle column:** Fully automated vs. manual extraction. **Right col-***

***umn:*** *Semi-automated vs. manual extraction. Blue dashed line: median difference (bias); red dashed lines: limits of agreement (LoA).*

D. Bland-Altman plots of the second-rater

Non-parametric Bland-Altman plots of 1160 data points (290 images *4 electrodes (Anode, Cathode$_{1-3}$)) from the second rater shows similar results compared with the first rater (**Figure 2**). For each pairwise method comparison across all montages and electrodes are shown in **Figure D.1**. The comparison between fully and semi -automated methods revealed the smallest bias and narrowest LoAs (bias = 1.16 mm; LoA: [−1.23, 3.48]), with 89.91% of data points (1043/1160) falling within the LoA (**Figure D.1A**). In contrast, comparisons involving manual extraction exhibited larger biases and greater variability. The fully automated vs. manual comparison showed a bias of −6.45 mm (LoA: [−10.69, −3.16]; **Figure D.1B**), while the semi-automated vs. manual comparison had a bias of −7.61 mm (LoA: [−12.75, −3.18]; **Figure D.1C**). Only 83.25% and 84.81% of points, respectively, fell within the LoA for these comparisons.

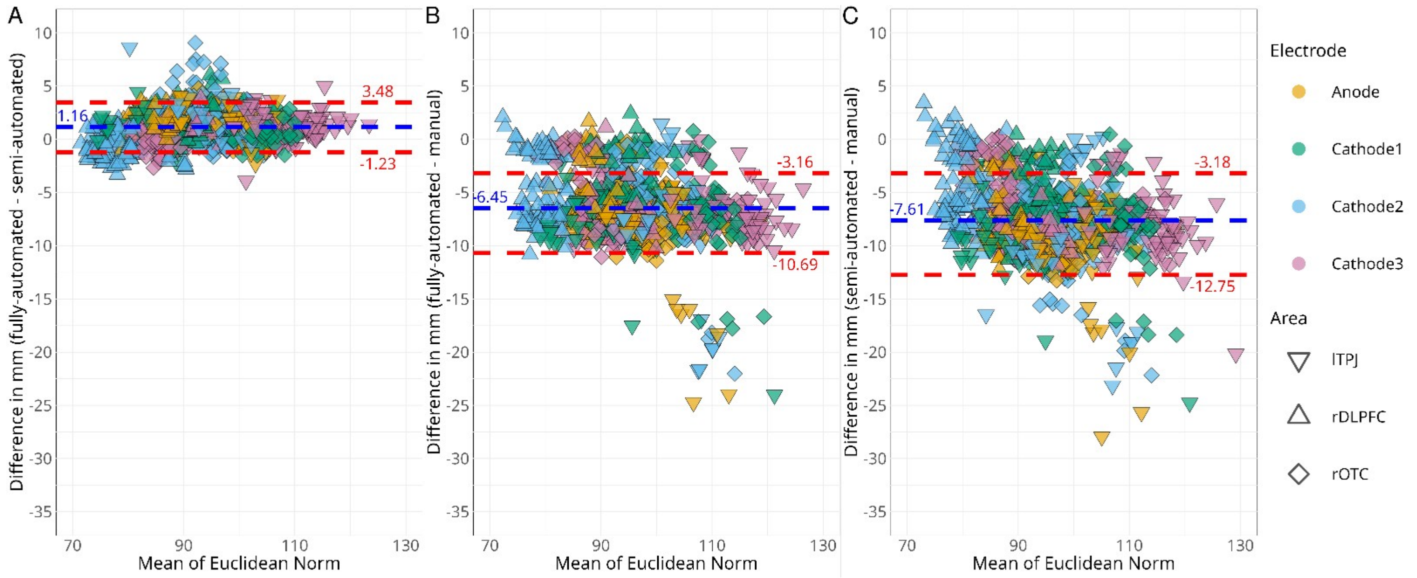


***Figure D.1****: Pairwise method comparison of the second-rater using Bland-Altman plots. Non-parametric Bland-Altman plots comparing electrode coordinate extraction methods across three montage areas. Differences (y-axis) between paired methods are plotted against their mean Euclidean distance (x-axis) for each electrode.* ***A)*** *Fully vs. semi-automated extraction.* ***B)*** *Fully automated vs. manual extraction.* ***C)*** *Semi-automated vs. manual extraction. Blue dashed line: median difference (bias); red dashed lines: limits of agreement (LoA). lTPJ (left temporoparietal junction), rDLPFC (right dorsolateral prefrontal cortex), rOTC (right orbito-temporal cortex)*

E. Normality test

| Test \ Diff | Diff (fully auto, semi-auto) | | Diff (fully auto, manual) | | Diff (semi-auto, manual) | |
|---|---|---|---|---|---|---|
| **Shapiro-Wilk** | W = 0.96 | $p < 0.001$ | W = 0.88 | $p < 0.001$ | W = 0.93 | $p < 0.001$ |
| **Kolmogorov-Smirnov** | D = 0.057 | $p < 0.001$ | D = 0.13 | $p < 0.001$ | D = 0.09 | $p < 0.001$ |

***Table E.1****: Normality test for method differences of Bland-Altman plots. 2 Tests metrics for Shapiro-Wilk test and the Kolmogorov-Smirnov test were calculated. Diff (Difference), auto (automated), W (W-statistic), D (D-statistic), p (p-value)*
